# Genetic association analyses highlight *IL6*, *ALPL*, and *NAV1* as three new susceptibility genes underlying calcific aortic valve stenosis

**DOI:** 10.1101/515494

**Authors:** Sébastien Thériault, Christian Dina, David Messika-Zeitoun, Solena Le Scouarnec, Romain Capoulade, Nathalie Gaudreault, Sidwell Rigade, Zhonglin Li, Floriane Simonet, Maxime Lamontagne, Marie-Annick Clavel, Benoit J. Arsenault, Anne-Sophie Boureau, Simon Lecointe, Estelle Baron, Stéphanie Bonnaud, Matilde Karakachoff, Eric Charpentier, Imen Fellah, Jean-Christian Roussel, Jean Philippe Verhoye, Christophe Baufreton, Vincent Probst, Ronan Roussel, the D.E.S.I.R. Study Group, Richard Redon, François Dagenais, Philippe Pibarot, Patrick Mathieu, Thierry Le Tourneau, Yohan Bossé, Jean-Jacques Schott

**Author notes:** These authors jointly supervised this work (Thierry Le Tourneau, Yohan Bossé, Jean-Jacques Schott). **Corresponding authors**: Yohan Bossé, Ph.D., Professor, Laval University, Department of Molecular Medicine, Canada Research Chair in Genomics of Heart and Lung Diseases, Institut universitaire de cardiologie et de pneumologie de Québec, Pavillon Marguerite-d'Youville, Y2106, 2725 chemin Sainte-Foy, Quebec City (Quebec), Canada, G1V 4G5; Jean-Jacques Schott, Ph.D., l’Institut du thorax, Unité Inserm UMR 1087 / CNRS UMR 6291, IRS-UN, 8 Quai Moncousu, BP 70721, 44007 Nantes cedex 1.

## Abstract

To date, only two replicated loci, *LPA* and *PALMD*, have been identified as causal genes for calcific aortic valve stenosis (CAVS) using genome-wide and transcriptome-wide association study (TWAS). To identify additional susceptibility genes for CAVS, we performed a GWAS meta-analysis totaling 5,115 cases and 354,072 controls of European descent. Four loci achieved genome-wide significance, including two new loci: *IL6* (interleukin 6) on 7p15.3 and *ALPL* (alkaline phosphatase) on 1p36.12. A TWAS integrating an eQTL study of 233 human aortic valves identified *NAV1* (neuron navigator 1) on 1q32.1 as a new candidate causal gene. The CAVS risk alleles were associated with higher mRNA expression of *NAV1* in valve tissues. Association results at the genome-wide scale showed genetic correlation with coronary artery disease and cardiovascular risk factors. Our study highlights three new loci implicating inflammation, mineralization and blood vessel integrity in CAVS pathogenesis and supports shared genetic etiology with cardiovascular traits.

## Main text

Calcific aortic valve stenosis (CAVS) is characterized by progressive thickening and calcification of the aortic valve leaflets causing blood flow obstruction, heart failure and death^1^. Symptomatic patients with CAVS are treated by surgical aortic valve replacement. CAVS is the second most frequent indication for cardiac surgery after coronary artery bypass grafting^2^. No medical treatment is available and conventional cardiovascular drugs tested in clinical trials of CAVS have failed to slow the progression of the disease and/or reduce associated adverse events^1,3-5^. The pathophysiological mechanisms associated with the development of CAVS remain elusive. Previous epidemiological studies on CAVS described familial patterns compatible with genetic inheritance and indicated that genetics contribute more than environmental factors to disease susceptibility^6,7^. Unbiased genomic approaches are just starting to elucidate the genetic determinants of CAVS^8-10^. To date, only three susceptibility loci have been associated with aortic valve calcification and/or stenosis in recent GWAS/TWAS^8-10^. The first GWAS identified *LPA*^9^, which encodes for lipoprotein(a), and our recent efforts using a TWAS approach identified *PALMD* as a candidate target gene for CAVS^8^. Another independent group reported associations at the *PALMD* and *TEX41* loci with aortic valve stenosis and bicuspid aortic valve^10^. The discovery of new loci to understand the genetic architecture of CAVS will provide invaluable information about key molecular processes and could open novel therapeutic avenues.

We undertook a GWAS meta-analysis including four cohorts: QUEBEC-CAVS, CAVS-France-1, CAVS-France-2 and UK Biobank (Online Methods). A total of 5,115 cases and 354,072 controls of European ancestry were included after quality control filters were applied (**Table 1**). We used the Haplotype Reference Consortium^11^ as reference to impute common variants. We performed a fixed-effects meta-analysis using the summary statistics for the association between ~ 8 million SNPs and CAVS risk in each cohort. The quantile-quantile plot indicated no inflation of observed test statistics (**Supplementary Fig. 1**). Four loci reached genome-wide significance (*P*_*GWAS*_ <5×10^−8^), including two previously described loci, namely *LPA* on 6q25.3-q26 and *PALMD* on 1p21.2^8,9^ (**Fig. 1)**, and two new loci: the interleukin 6 (*IL6*) gene on 7p15.3 (rs2069832, *P*=1.1×10^−8^) and the alkaline phosphatase (*ALPL*) gene (rs12141569, *P*=3.9×10^−8^) on 1p36.12 (**Fig. 1 & Fig. 2**). The lead SNPs for all loci with *P*_*GWAS*_ <1×10^−5^ are indicated in **Supplementary Table 1**.

**Figure 1.**
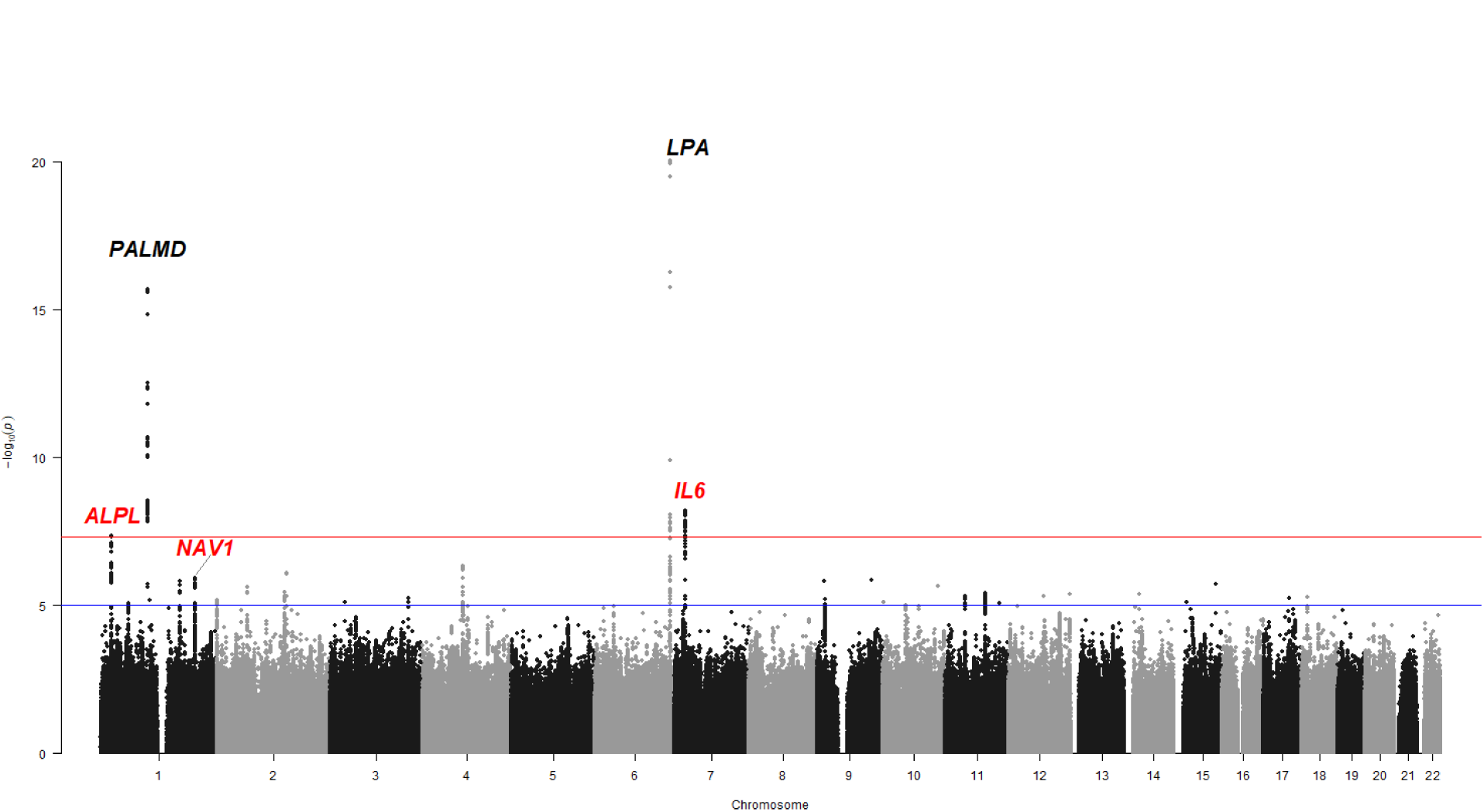
Manhattan plot of the GWAS meta-analysis of four cohorts including 5,115 cases and 354,072 controls. The y axis represents *P* value in −log10 scale. The horizontal red and blue lines indicate *P* values of 5×10^−8^ and 1×10^−5^, respectively. Gene names in black and red correspond to previously known and novel loci for CAVS, respectively.

**Figure 2.**
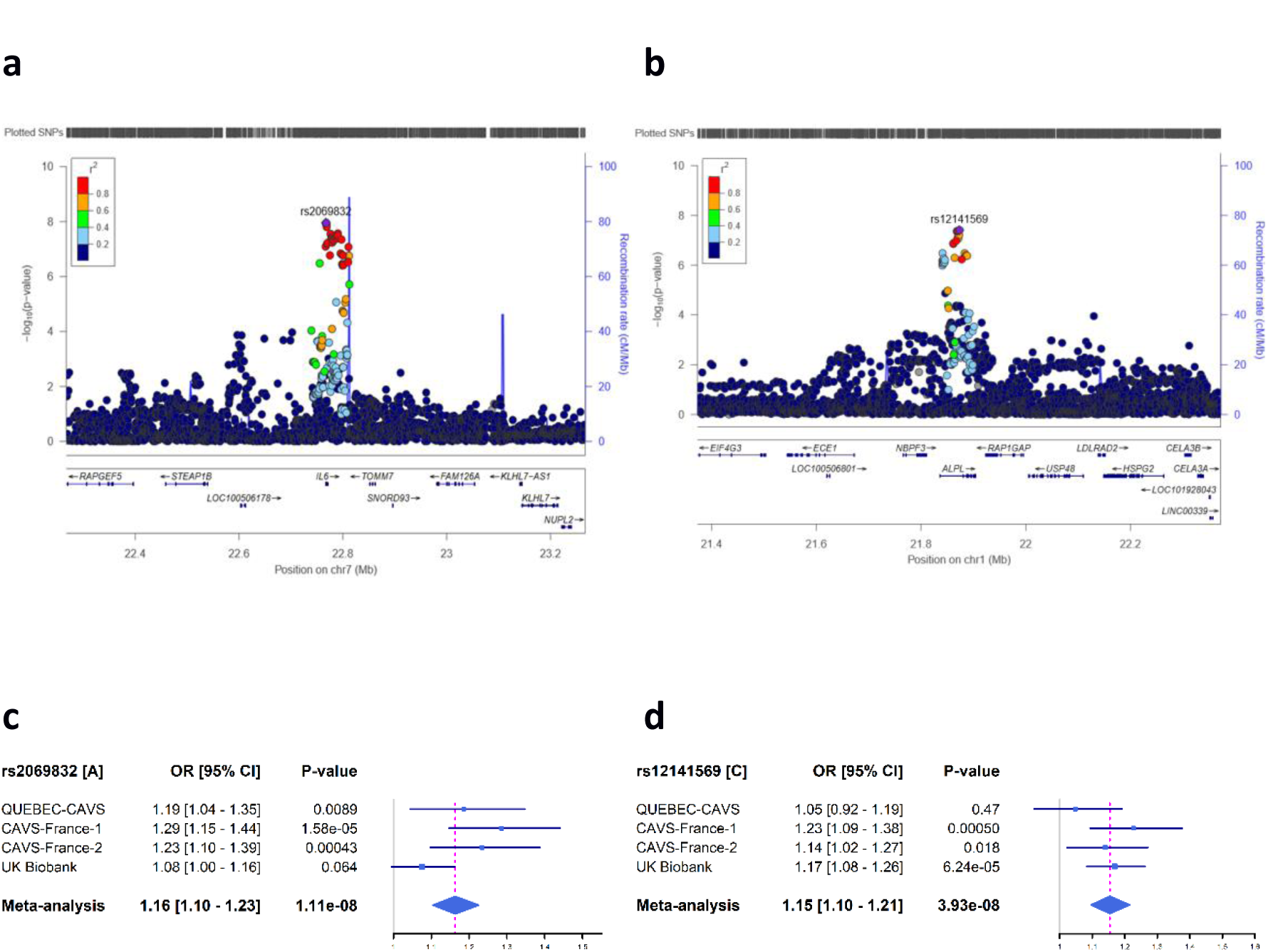
Two new CAVS-associated loci. (**a-b**) Regional plots showing the two new loci. The y axis shows the *P* values in −log10 scale for SNPs up- and downstream of the sentinel SNP (purple dot). The extent of linkage disequilibrium (LD; r^2^ values) for all SNPs with the sentinel SNP is indicated by colors. The location of genes is shown at the bottom. SNPs are plotted based on their chromosomal position on build hg19. (**c-d**) Forest plots showing the effect size of the top CAVS-associated SNP at the two new loci in each cohort and meta-analysis. The blue filled squares represent the odds ratio (OR) for each cohort. The horizontal lines represent the 95% confidence intervals of the OR. The grey and the dashed magenta vertical lines represent an OR of 1.0 and the OR of the meta-analysis, respectively.

**Table 1.** Sample size for the GWAS meta-analysis on CAVS

| **Cohort** | **Cases (n)** | **Controls (n)** |
| --- | --- | --- |
| QUEBEC-CAVS | 1,009 | 1,017 |
| CAVS-France-1 | 1,261 | 1,305 |
| CAVS-France-2 | 1,495 | 2,707 |
| UK Biobank | 1,350 | 349,043 |
| **Total** | **5,115** | **354,072** |

We have then integrated the summary statistics from the four-cohort meta-analysis with our valve eQTL dataset (n=233) to perform a new TWAS on CAVS. *PALMD*, identified in our previous TWAS^8^, is strongly associated with CAVS (*P*_*TWAS*_ =3.6×10^−17^; **Fig. 3a**). After correction for the number of genes with significant *cis*-heritability to calculate expression weights in the valve eQTL dataset (n=11,834), one additional gene reached the significance threshold (*P*_*TWAS*_ <4.2×10^−6^), namely neuron navigator 1 (*NAV1*) (*P*_*TWAS*_ =2.0×10^−6^). *NAV1* is located on chromosome 1q32.1; the lead GWAS SNP is rs665770, located in intron 3 (*P*_*GWAS*_ =1.5×10^−6^) (**Supplementary Fig. 2 and Fig. 3b**). It is also the lead valve eQTL-SNP (*P*_*eQTL*_ =5.9×10^−11^). The risk allele for CAVS (“A”) is associated with higher mRNA expression levels of *NAV1* in valve tissues (**Fig. 3c**). Moreover, Bayesian colocalization test^12^ indicated a high probability of shared GWAS and *NAV1* eQTL signals (PP4=98.3%). Finally, the CAVS risk alleles were consistently associated with increased expression of *NAV1* in aortic valve tissues (**Fig. 3d**). Together, these results suggest that this locus confers susceptibility through higher expression of *NAV1* in the aortic valve.

**Figure 3.**
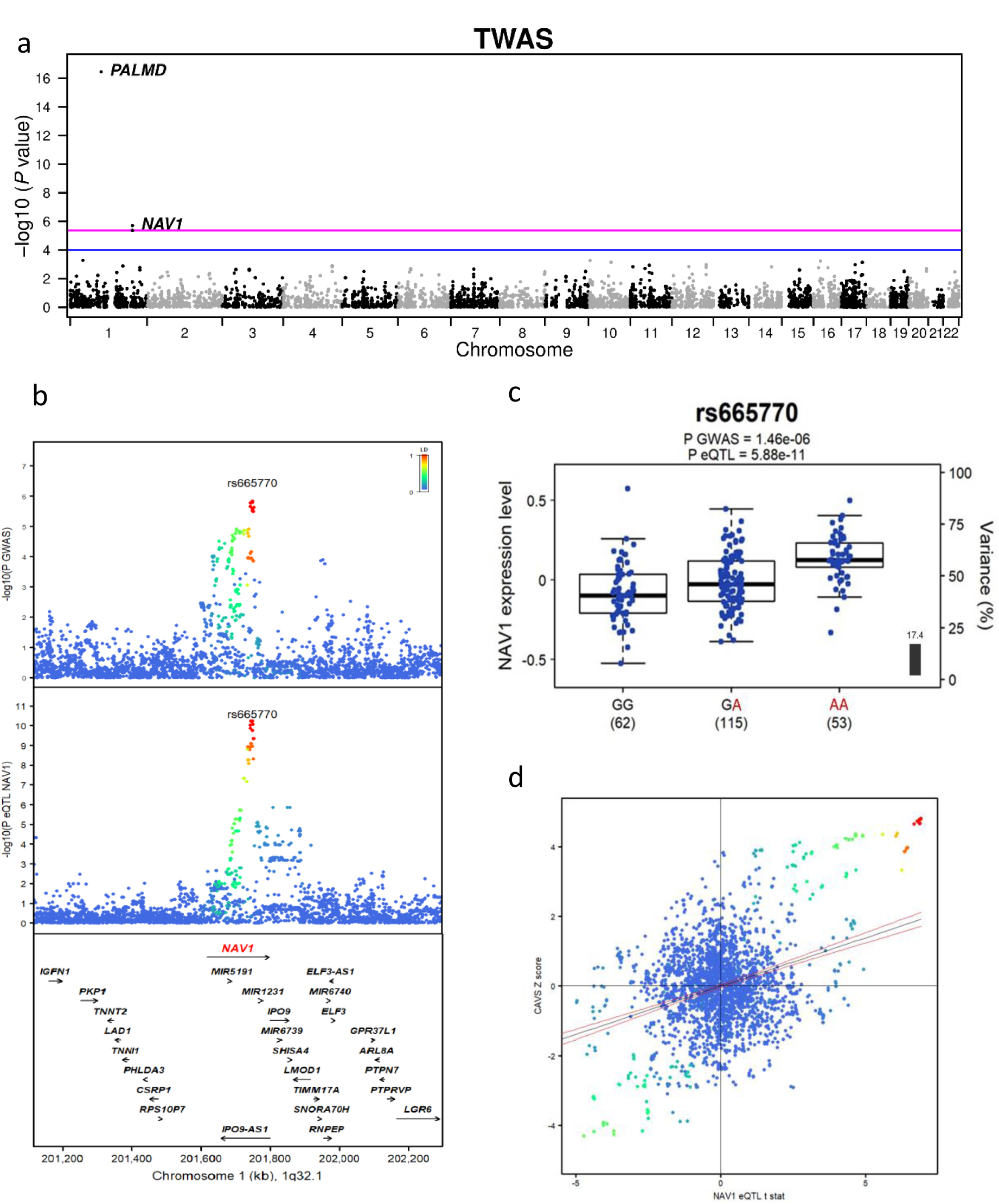
Identification of *NAV1* by TWAS and causality analysis. (**a**) TWAS in valve tissue with CAVS. *P* values for gene expression-CAVS associations are on the y-axis in −log10 scale. The blue and magenta horizontal lines represent *P*_*TWAS*_ of 0.0001 and 4.23 x 10^−6^ (Bonferroni), respectively. Annotations for the top significant probes are indicated. (**b**) GWAS and valve eQTL results surrounding *NAV1* on chromosome 1q32.1. The upper panel shows the genetic associations with CAVS. The bottom panel shows the valve eQTL statistics for the *NAV1* gene. The extent of linkage disequilibrium (LD; r^2^ values) for all SNPs with rs665770 is indicated by colors. The location of genes in this locus is illustrated at the bottom. (**c**) Boxplots of gene expression levels in the valves according to the three genotype groups for SNP rs665770. The y-axis shows the mRNA expression levels. The x-axis represents the three genotype groups for rs665770 with the number of individuals in parenthesis. Expression data for the *NAV1* probe was not available for 3 individuals (n=230). The risk allele identified in our GWAS meta-analysis is shown in red. Box boundaries, whiskers and center marks in boxplots represent the first and third quartiles, the most extreme data point which is no more than 1.5 times the interquartile range, and the median, respectively. The black bar and the left y axis indicate the variance in *NAV1* gene expression explained by rs665770. (**d**) Scatterplot of the 1q32.1 susceptibility locus showing SNP associations with CAVS and *NAV1* gene expression in aortic valve tissues. The y axis represents variant association with CAVS (Z score). The x axis shows association with *NAV1* gene expression (t statistic). Variants are colored based on the degree of LD (r^2^) with the top CAVS-associated variant rs665770. The blue line is the regression slope with 95% confidence interval (red lines).

Overall, three new CAVS risk loci were discovered in this study: *IL6*, *ALPL*, and *NAV1*, among which only *NAV1* had significant eQTL in our aortic valve dataset. A series of post-GWAS analyses were then carried out to determine the variants most likely to be functional and gain further insight about these new CAVS loci. We also sought to identify pathways, cell or tissue types driving the genetic risk of CAVS and assessed the genetic correlation of CAVS with other traits/diseases.

Fine-mapping to localize the potential causal variants at CAVS loci was performed using PAINTOR^13^ with the following annotations: conservation score, fetal DNase I hypersensitive sites (DHS), promoter region and H3K4me1 peaks. For our previously reported *PALMD* locus, the lead GWAS SNP, rs6702619, provides a posterior probability for causality (PPC) of 0.98. For the *IL6* locus, rs1800795 achieved a similarly strong PPC of 0.96. This SNP is located <1 kb from the lead SNP rs2069832 (LD r^2^=0.97 in 1000 Genomes Europeans). For the *ALPL* locus, the index SNP, rs12141569, has the top PPC of 0.55, but a second plausible functional variant was identified (rs7547128 with a PPC of 0.21). Finally, three SNPs were prioritized for the *NAV1* locus, including rs7535989, rs665770 (GWAS lead SNP), and rs665834 with PPCs of 0.54, 0.15, and 0.06, respectively. Predicted effects on chromatin features using the DeepSEA^14^ algorithm for the lead SNPs and variants identified using PAINTOR at the three new CAVS loci are available in **Supplementary Table 2**.

CAVS-associated variants were evaluated for eQTL in GTEx^15^. rs2069832-*IL6* is a strong eQTL in many tissues (adipose tissues, arteries, breast, esophagus, heart, lung, and others) for the *IL6* RNA antisense (LOC541472) (**Supplementary Fig. 3**). The direction of effect is consistent across all tissues, i.e. higher expression corresponds to higher risk of CAVS. Other eQTLs of note in GTEx include rs12141569-*ALPL* in fibroblasts (*P*_*eQTL*_ =1.1×10^−7^) and rs665770-*NAV1* in heart atrial appendage (*P*_*eQTL*_ =1.9×10^−8^). The lead SNPs at the three new loci were further investigated using a phenome-wide association study (PheWAS) in 353,378 individuals of European ancestry from the UK Biobank, testing a total of 832 curated phenotypes. The *IL6* variant rs2069832 was strongly associated with eosinophil count (*P*=7.9×10^−24^) and to a lower extent with leukocyte, lymphocyte and neutrophil counts, asthma, systolic blood pressure, and carotid artery procedures (**Fig. 4a**). The CAVS risk allele was associated with higher eosinophil count, systolic blood pressure and carotid artery procedures, but surprisingly lower risk of asthma. For *ALPL*, no phenotype was associated with rs12141569 after correction for multiple testing, but a positive nominal association was observed between the CAVS risk allele and bone mineral density (*P*=4.7×10^−4^, **Fig. 4b**). For *NAV1*, the CAVS risk allele for rs665770 was associated with lower diastolic blood pressure (*P*=6.5×10^−10^) as well as higher risk of carotid artery stenosis (*P*=7.4×10^−8^) and carotid artery procedures (*P*=1.7×10^−5^) (**Fig. 4c**). Bayesian colocalization^12^ demonstrated shared GWAS signals between our CAVS meta-analysis, carotid stenosis (PP4=98.5%) and carotid procedures (PP4=97%) in UK Biobank (**Supplementary Fig. 4**).

**Figure 4.**
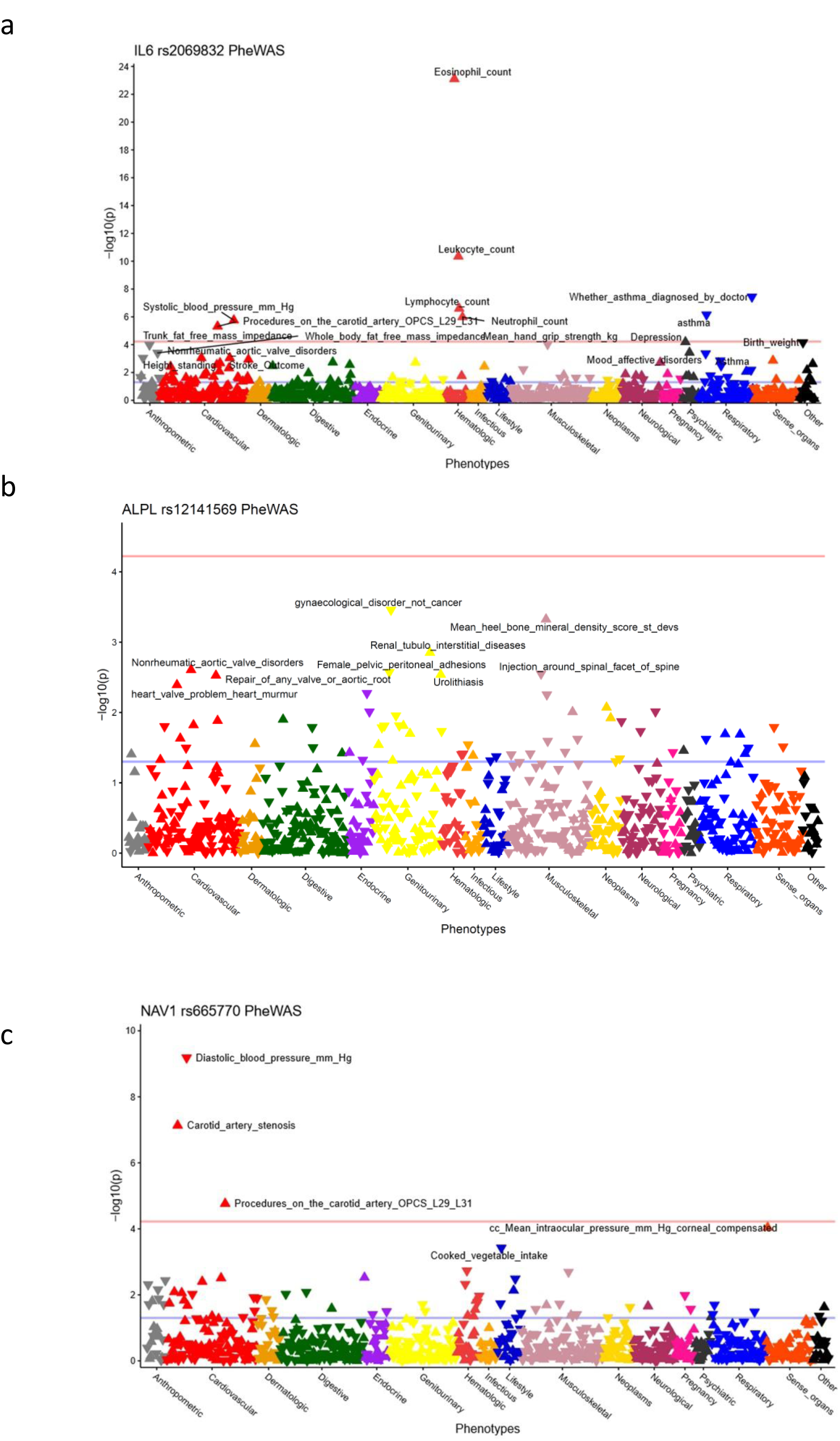
PheWAS for the three new CAVS risk loci in UK Biobank. Each triangle represents a different phenotype (n=832). Triangles pointing up and down are positive and negative associations with CAVS risk alleles, respectively. The pink horizontal line represents the threshold for significance after correcting for multiple testing (p=0.05/832=6.0×10^−5^). The blue horizontal line represents the threshold for nominal significance (p=0.05).

Although the exact pathophysiological mechanisms remain to be elucidated, the three identified genes can be functionally linked to CAVS. Interleukin 6 has been identified previously as a strong promoter of valve interstitial cells mineralization^16^. The RNA antisense modulated in various tissues by our lead variant is strongly upregulated in monocytes during fungal infection^17^ and was shown to induce *IL6* expression in glioma cells by causing the enrichment of histone H3 acetylated at lysine 27 (H3K27Ac) of the *IL6* promoter^18^. The *ALPL* gene codes for tissue non-specific alkaline phosphatase (ALPL, also called TNAP), a crucial enzyme involved in the mineralization process^19^, and was found to be enriched in calcified aortic valves as compared to non-calcified valves^20^. Although our lead variant was not associated with blood alkaline phosphatase levels in previous GWAS^21-23^, an effect on ALPL activity is possible, as suggested from our PheWAS analysis as well as a recent GWAS showing an even stronger association of rs12141569 with bone mineral density (*P*=9.9×10^−9^)^24^. *NAV1* is strongly expressed in the human heart^25^, as well as in the aorta (GTEx Portal); our lead SNP shows eQTLs in aortic valve tissue which colocalizes with CAVS risk as well as with carotid stenosis in UK Biobank. This locus was also associated with diastolic blood pressure and pulse pressure in previous GWAS^26,27^, pointing to an effect on blood vessel integrity. Arterial blood pressure and the aortic root morphology have been previously identified as risk factors for CAVS and its progression over time^28,29^.

Tissue and gene-set enrichment were performed with DEPICT^30^ using SNPs nominally associated with CAVS (*P*_*GWAS*_ <1×10^−5^). At the tissue level, enrichments with nominal *P*_*DEPICT*_ <0.05 were observed for arteries, veins and blood vessels, but none were significant at FDR<5%. Similarly, no gene sets were significantly enriched at FDR<5%. Top nominal pathways of interest include increased compact bone thickness, increased heart weight, disorganized myocardium and increased systemic arterial diastolic blood pressure (**Supplementary Table 3)**. We also tested enrichment of CAVS-associated loci (*P*_*GWAS*_ <1×10^−5^) in DHS in cell lines and tissues from ENCODE and the Epigenome Roadmap Project using FORGE^31^ and GARFIELD^32^. Leading tissues were heart, blood vessels and fibroblast, but no overlap reached statistical significance (**Supplementary Fig. 5**). LD score regression^33^ was then used to test genetic correlation with other traits and diseases (**Supplementary Table 3)**. We discovered significant genetic correlation with CAD, myocardial infarction, diastolic and systolic blood pressure, body mass index, type 2 diabetes and large-artery stroke (**Table 2**).

**Table 2.** Genetic correlation between CAVS and other traits and diseases by LD score regression

| **Traits/diseases** | **Correlation (Rg)** | **P** |
| --- | --- | --- |
| Coronary artery disease | 0.23 | 0.0001 |
| Myocardial infarction | 0.26 | 0.0004 |
| Systolic blood pressure | 0.15 | 7.74x 10<sup>-8</sup> |
| Diastolic blood pressure | 0.12 | 1.90x10<sup>-5</sup> |
| BMI | 0.21 | 6.41x10<sup>-6</sup> |
| WHR | 0.20 | 0.0010 |
| WHR adjusted for BMI | 0.08 | 0.1798 |
| Type 2 diabetes | 0.18 | 6.08x10<sup>-6</sup> |
| Total cholesterol | 0.12 | 0.0247 |
| LDL-cholesterol | 0.14 | 0.0125 |
| HDL-cholesterol | -0.14 | 0.0073 |
| Triglyceride | 0.16 | 0.0070 |
| Any ischemic stroke | 0.16 | 0.0011 |
| Large-artery stroke | 0.20 | 2.00x10<sup>-5</sup> |
| Cardioembolic stroke | 0.15 | 0.0024 |
| Small-vessel stroke | 0.07 | 0.2159 |
| Chronic kidney disease | 0.10 | 0.1727 |

BMI: Body-mass index; WHR: Waist-to-hip ratio; LDL: Low-density lipoprotein; HDL: High-density lipoprotein.

In conclusion, we describe three new risk loci for CAVS (*IL6*, *ALPL* and *NAV1*) implicating inflammation, mineralization and blood vessel integrity in the disease pathogenesis. These genes provide key insights into the etiology of CAVS and constitute novel targets for the development of therapeutic agents. Our association results at the genome-wide scale show genetic correlation with cardiovascular disease and cardiovascular risk factors, supporting a shared genetic etiology.

## Methods

### Study cohorts

#### QUEBEC-CAVS

A complete description of the cohort has already been published^8^. Briefly, blood samples and aortic valves were collected from patients with severe aortic valve stenosis undergoing aortic valve replacement at the Institut universitaire de cardiologie et de pneumologie de Québec. Only cases with tricuspid nonrheumatic CAVS were included. No other severe valvular heart diseases were present. CAVS cases underwent a comprehensive Doppler echocardiographic examination; the transvalvular gradient was calculated using the modified Bernoulli equation and the aortic valve area calculated with the continuity equation. In parallel, a control group matched for ethnicity, age, gender, type 2 diabetes and hypertension was recruited from patients that underwent cardiac surgery, mostly for isolated coronary artery bypass (>98%). Absence of CAVS was confirmed by echocardiography. All patients signed an informed consent for the realization of genetic studies. The study was approved by the ethics committee of the Institut universitaire de cardiologie et de pneumologie de Québec. Coronary artery disease was defined as a history of myocardial infarction, coronary artery stenosis on coronary angiography, or documented myocardial ischemia.

Blood samples were collected and DNA was extracted from frozen buffy coat. Whole-genome genotyping was performed using the Illumina HumanOmniExpress BeadChip. Standard genotyping quality control (QC) procedures were performed^8^. After the QC filters, 1,009 cases and 1,017 controls were available for subsequent analyses. Genotypes were imputed with the Michigan Imputation Server^34^ using the Haplotype Reference Consortium version 1 (HRC.r1-1) data^11^ as reference set.

Genetic association tests were performed using additive logistic regression models based on expected genotype counts (dosages) as implemented in the software SNPTEST v2.5.2^35^, adjusting for age, sex, and the first 10 ancestry-based principal components.

### French Datasets

A large cohort of “isolated” CAVS cases has been constituted by l’institut du thorax in Nantes. A Doppler-echocardiography and blood sampling were carried out at the time of enrollment. Patients with severe renal failure, history of rheumatic disease or chest radiation were excluded. So far, 1,663 severe CAVS cases (including patients with tricuspid or bicuspid valves), have been recruited from 2001 to 2017 thanks to a collaboration between Nantes, Rennes and Angers University Hospitals. Most of them have been referred to surgery after enrollment and valve tissue was collected in a subset of patients. The study was approved by the local ethics committee and all patients signed an informed consent for the purpose of genetic studies. Coronary artery disease was defined as previously described in the QUEBEC-CAVS cohort.

In parallel, the Cardiovascular department in Bichat university Hospital in Paris has recruited 1,500 patients with CAVS, including patients with tricuspid or bicuspid valves (GENERAC and COFRASA projects). All patients were prospectively enrolled and fully phenotyped. Blood samples and valve tissues were collected (DNA, blood and tissue bank stored at the level of our Center of Biological Resources).

The control populations came from two datasets called D.E.S.I.R. and PREGO. D.E.S.I.R. (The Data from the Epidemiological Study on the Insulin Resistance Syndrome)^36,37^ is an epidemiological cohort used here as a control general population. PREGO (Population de Référence du Grand Ouest) is a set of 5,707 healthy persons selected through the Blood Donor Service, originating from Western France, as a resource dedicated to provide a regional reference population of Western France for national and international research projects in the field of evolution, population and medical genetics. CAVS status was not available in D.E.S.I.R. and PREGO because of the lack of echocardiography data in this cohort.

The patients were genotyped in two waves:

### CAVS-France-1

A set of 1,329 patients with CAVS from the *institut du thorax* biobank were genotyped using Axiom Genome-Wide CEU-1 array (Affymetrix, Inc). We used a general population as controls. A subset of 901 individuals came from the D.E.S.I.R. population and 466 from the PREGO population.

We applied quality control procedures to filter markers (genotyping rate and heterozygosity) and individuals (relatedness and demographic stratification). The ancestry of participants was assessed using a multi-dimensional scaling technique implemented in PLINK^38^. Multi-dimensional scaling method was applied on the Identity-By-State matrix using cases and controls together with all 1000G populations and all non-Finnish European populations. We excluded outliers on the first two components using an expectation-maximization (EM)-fitted Gaussian mixture clustering method implemented in the R package M-CLUST^39^, assuming one cluster and noise. Outlier position was initialized using a nearest-neighbour-based classification (NNclust in R package PrabClus)^40^.

After quality control, we kept 1,261 CAVS patients (18.5% had a bicuspid valve) and 1,305 controls (865 individuals from the D.E.S.I.R. population and 440 from the PREGO population). A final list of 350,488 SNPs were used for imputation (138,199 SNPs were removed for genotyping rate or low MAF or HWE deviation and 1,095 SNPs removed due to technical differences between genotyping plates).

### CAVS-France-2

The French CAVS-France-2 dataset was genotyped using Axiom Genome-Wide PMRA array (Affymetrix, Inc). The dataset is composed of a set of 1,478 patients recruited at the Hôpital Bichat, 319 patients from the *institut du thorax* biobank and 2,828 controls from the PREGO population.

We applied the same quality control procedures and selection on individuals as for the CAVS-France-1 dataset. 1,181 patients from Hôpital Bichat, 314 patients from the *institut du thorax* biobank and 2,707 controls remained for analysis. Among the CAVS cases, 23.9% had a bicuspid valve. A final list of 207,518 SNPs were used for imputation (275,504 SNPs were removed for genotyping rate or low MAF or HWE deviation and 576 SNPs removed due to technical differences between genotyping plates).

Imputation for both France cohorts was performed using the Michigan Imputation Server^34^ with the Haplotype Reference Consortium version 1 (HRC.r1-1) data^11^ as reference set.

Genetic association tests were performed using additive logistic regression models based on expected genotype counts (dosages) as implemented in the software SNPTEST v2.5.2^35^, adjusting for the first 5 ancestry-based principal components as well as the components associated with the disease among the fifth to tenth principal components.

### UK Biobank

UK Biobank is a large prospective cohort of about 500,000 individuals between 40 and 69 years old recruited from 2006 to 2010 in several centers located in the United Kingdom^41^. The present analyses were conducted under UK Biobank data application number 25205. CAVS diagnosis was established from hospital record, using the International Classification of Diseases version-10 (ICD10) and Office of Population Censuses and Surveys Classification of Interventions and Procedures (OPCS-4) coding. CAVS was defined as ICD10 code number I35.0 or I35.2.

Participants with a history of rheumatic fever or rheumatic heart disease as determined by ICD10 codes I00–I02 and I05–I09 were excluded from the CAVS group. We included all other participants in the control group, except for those with OPCS-4 codes K26 or K30.2 or a self-reported diagnosis of CAVS, which were excluded from the analysis. Samples were genotyped with the Affymetrix UK BiLEVE Axiom array or the Affymetrix UK Biobank Axiom Array. Phasing and imputation were performed centrally using the Haplotype Reference Consortium (HRC) reference panel^11^. Samples with call rate <95%, outlier heterozygosity rate, gender mismatch, non-white British ancestry, related samples (second degree or closer), samples with excess third-degree relatives (>10), or not used for relatedness calculation were excluded.

Variants not on both arrays, which failed QC in more than one batch, with call rate <95% or with minor allele frequency (MAF) <0.01 were excluded. To estimate the association of each variant with CAVS, we performed linear regression following an additive model using BGENIE (BioRxiv: https://www.biorxiv.org/content/early/2017/07/20/166298), with adjustment for age, sex and the first 20 ancestry-based principal components. This program was developed to perform efficient GWAS in large populations and was specifically written for the analysis of the UK Biobank dataset.

### Meta-analysis

The genomic inflation factors in the four cohorts were 1.029 (QUEBEC-CAVS), 1.031 (CAVS-France-1), 1.050 (CAVS-France-2), and 1.021 (UK Biobank). Genomic correction was applied to each cohort. Variants with an r^2^ value of ≤0.3 or MAF <0.01 were removed in each cohort. We performed a fixed-effects sample size-weighted meta-analysis as implemented in METAL^42^. The effective sample size was calculated as 4/(1/N_cases_ + 1/N_controls_) to account for imbalance in the number of cases and controls in each cohort. The weights were proportional to the square-root of the effective sample size. Genomic correction was applied to the results of the meta-analysis (lambda =1.022). For variants associated with CAVS below a *P* value threshold of 1×10^−5^ in the sample size weighted meta-analysis, we used logistic regression models in UK Biobank, as implemented in the software SNPTEST v2.5.2^35^, to obtain effect size estimates. Models were based on expected genotype counts (dosages), with adjustment for age, sex, and the first 10 ancestry-based principal components. We then performed the meta-analysis again for these variants using a fixed-effects inverse-variance weighted method. Heterogeneity was evaluated using Cochran’s Q-test as implemented in METAL^42^. The genome-wide significant *P* value cutoff was set to 5×10^−8^.

### Transcriptome-wide association study (TWAS)

The TWAS was performed using FUSION^43^. Our aortic valve eQTL dataset, which includes 233 individuals, was used as reference^8^. Gene expression weights were calculated using prediction models implemented in FUSION. This includes top1 (i.e. the single most significant valve eQTL-SNP as the predictor), LASSO regression, and enet (elastic net regression). SNP data located 500 kb on both sides of the probes were used to obtain expression weights. All probes that passed QC in the valve eQTL were evaluated (n=45,699). There were 11,384 genes with significant *cis*-heritability (*P*<0.01) to calculate expression weights. Expression weights were then combined with summary-level results from the meta-analysis to estimate association statistics between gene expression and CAVS. Genome-wide significant TWAS genes were considered at *P*_TWAS_<4.2×10^−6^ (Bonferroni =0.05/11,384).

### Post-GWAS analyses

#### Fine-mapping to identify causal variants

We used Probabilistic Annotation INTegratOR (PAINTOR)^13^ to identify candidate causal variants. This program uses a probabilistic framework to integrate LD information, *P* value distribution in significant loci and functional annotations. We used all SNPs located within 500 kb of the genome-wide significant loci (*LPA*, *PALMD*, *ALPL*, *IL6* and *NAV1*) and with a *P* value <0.01 for association with CAVS in the GWAS meta-analysis. LD structure was obtained from the 1000 Genomes European reference set. As described previously^44^, we used genetic annotations that were not cell-type specific. The recommended number of annotations to use in PAINTOR is 3 to 5. After determining the significance of each annotation, we selected the following 4 annotations to optimize the model: conservation score, fetal DNase I hypersensitive sites, promoter region and H3K4me1 peaks. Additional annotations did not increase the model-fit or were highly correlated with the 4 selected annotations. We used a model allowing for 1 causal variant.

#### Investigation of transcriptomic and epigenetic effects

We looked for eQTL associations for the lead SNPs at the newly identified loci in the Genotype-Tissue Expression (GTEx-V7) project data^15^. This resource provides eQTL data from 48 tissues (620 individuals).

We looked at the effects of the variants with suggestive association with CAVS in the meta-analysis (*P* value <1×10^−5^) on transcription factor binding and chromatin marks in 149 cell lines from ENCODE and the Roadmap Epigenomics project using DeepSEA^14^.

#### Colocalization

When the newly identified loci had significant eQTL associations, we tested for colocalization between the CAVS risk association and the aortic valve or other tissues eQTL association. We used the COLOC package in R to perform Bayesian colocalization analyses^12^. COLOC tests for five hypotheses: H0, no eQTL and no GWAS association; H1, association with eQTL, but no GWAS; H2, association with GWAS, but no eQTL; H3, eQTL and GWAS associations, but independent signals; and H4, shared eQTL and GWAS associations. In practice, a high posterior probability of H4 (PP4>75%) indicates colocalization.

#### Phenome-wide association studies

We verified the association between the lead variant in the two new genome-wide significant loci (*ALPL* and *IL6*) as well as the new locus identified with TWAS (*NAV1*) and a wide range of phenotypes in the UK Biobank. We curated 832 phenotypes from anthropometric traits, health questionnaires, ICD10 diagnostic codes, OPCS-4 procedure codes, imaging data and laboratory markers. Phenotypes were grouped in the following categories: anthropometric, cardiovascular, dermatologic, digestive, endocrine, genitourinary, hematologic, infectious, lifestyle, musculoskeletal, neoplasms, neurological, pregnancy, psychiatric, respiratory, sense organs and others. Continuous variables were examined individually to exclude outlier values and quantile normalization was applied. For each phenotype, an additive logistic (binary phenotype) or linear regression (continuous phenotype) test was performed in 353,378 unrelated individuals of White-British ancestry, adjusting for age, sex and the first 10 ancestry-based principal components using SNPTEST v2.5.2^35^. Results for each phenotype were plotted using the PheWAS R package. A significance threshold of *P*=6.0×10^−5^ (0.05/832) was applied to correct for multiple testing. We used the colocalization procedure described above to test for shared causal association between CAVS and other phenotypes identified in the PheWAS.

#### Pathway analyses and tissue enrichment

We used data-driven expression-prioritized integration for complex traits (DEPICT)^30^ to identify over-represented biological pathways and enrichment in tissues among the variants with suggestive association with CAVS (P<1×10^−5^). We selected independent variants by applying LD pruning based on European individuals from the 1000 Genomes using a threshold of r^2^<0.05. We tested enrichment of CAVS-associated loci (P<1×10^−5^) in DNase I sensitivity hotspots in cell lines and tissues from ENCODE and the Roadmap Epigenomics project using the Functional element Overlap analysis of the Results of Genome Wide Association Study Experiments (FORGE) tool^31^. We used GARFIELD^32^ to correlate our GWAS findings with regulatory or functional annotations and find features relevant to a phenotype of interest. Since our GWAS included only European individuals, we used the original files describing the allele frequencies and linkage disequilibrium from the UK10K data provided in the GARFIELD distribution. We also used the annotation and distance to TSS files provided with the GARFIELD release. The annotations included 1,005 features extracted from ENCODE, GENCODE and Roadmap Epigenomics projects, including genic annotations, chromatin states, histone modifications, DNase I hypersensitive sites and transcription factor binding sites, among others, in a number of publicly available cell lines.

#### Genetic correlation

We performed LD-score regression to estimate the genetic correlation between CAVS and other cardiovascular traits^33,45^. We used the results of the sample size weighted CAVS GWAS meta-analysis (performed without genomic correction). Summary statistics from GWAS meta-analyses on cardiovascular traits (coronary artery disease, blood pressure, hypertension, blood lipids, type 2 diabetes, body mass index, waist-to-hip ratio, stroke, chronic kidney disease) were obtained from publicly available resources (**Supplementary Table 4**). For each trait, we removed SNPs located in the major histocompatibility complex region (chr 6; positions 28,477,797–33,448,354 GRCh37), SNPs with an extreme effect size (chi^2^ statistic >80) and selected a subset of SNPs included in HapMap3 to optimize imputation quality. We used the European LD-score files calculated from 1000 Genomes reference panel provided by the developers. For the blood pressure traits, since both populations included the UK Biobank, we constrained the intercept based on the amount of overlap and correlation between CAVS and the trait evaluated. Analyses were performed using the ldsc program (https://github.com/bulik/ldsc/). A conservative *P* value threshold of <0.001 was considered significant.

#### Statistical analysis

Statistical analyses were performed with R version 3.2.3 unless otherwise specified. Two-sided *P* values below 0.05 were considered significant unless otherwise specified.

## Acknowledgements

We thank the research team at the cardiac surgical database and biobank of the Institut universitaire de cardiologie et de pneumologie de Québec (IUCPQ) for their valuable assistance. This research has been conducted using the UK Biobank Resource. S.T. holds a Junior 1 Clinical Research Scholar award from the Fonds de Recherche du Québec-Santé (FRQS). M.-A.C. holds a Junior 1 Research Scholar award from the FRQS. B.J.A. holds a Junior 2 Research Scholar award from the FRQS. P.P. holds the Canada Research Chair in Valvular Heart Disease and his research program is supported by a Foundation Scheme Grant from Canadian Institutes of Health Research (Ottawa, Ontario, Canada). P.M. holds a Fonds de Recherche du Québec-Santé (FRQS) Research Chair on the Pathobiology of Calcific Aortic Valve Disease. Y.B. holds a Canada Research Chair in Genomics of Heart and Lung Diseases. This work was supported by the Heart and Stroke Foundation of Canada and the Canadian Institutes of Health Research [MOP - 102481, MOP – 137058, PJT – 153396, PJT – 159641] to Y.B.

We thank Marie Marrec and Guénola Coste for their contribution to clinical data collection. We are most grateful to the Genomics and Bioinformatics Core Facility of Nantes (GenoBiRD, Biogenouest) for its technical support. This work was supported by an ANR & FRM grant [13-BSV6-0011, DCV20070409278] to JJS, by a Fédération Française de Cardiologie, a Fondation Coeur et Recherche and an Inserm Translational Research grant to T.L.T., a PHRC Interregional (API20-20) grant to V.P., a “Connect Talent” research chair from Région Pays de la Loire and Nantes Métropole to R.C. and the French Regional Council of Pays-de-la-Loire (VaCaRMe program) to R.Re.

We would like to specially thank the team of the Centre d'Investigation Clinique (Xavier Duval), the Centre de Ressources Biologiques (Sarah Tubiana), Christophe Aucan from the Assistance Publique - Hopitaux de Paris, Département de la Recherche Clinique et du Développement (DRCD) and Estelle Marcault from the Unité de Recherche Clinique Paris Nord for their help and support during all these years.

The PREGO bio-collection is part of the VaCaRMe program founded by the French Regional Council of Pays-de-la-Loire (RFI VaCaRMe). The authors wish to thank the Fondation Genavie for its financial support. We would like to specially thank Stephanie Chatel for all her contribution to constitute this bio-collection. We thank the Etablissement Français du Sang and the Centre de Ressources Biologiques of CHU de Nantes, for its support. We thank our numerous collaborators for the recruitment of blood donors, and all participating individuals.

The D.E.S.I.R. study has been supported by INSERM contracts with CNAMTS, Lilly, Novartis Pharma and Sanofi-Aventis; by INSERM (Réseaux en Santé Publique, Interactions entre les déterminants de la santé), Cohortes Santé TGIR, the Association Diabète Risque Vasculaire, the Fédération Française de Cardiologie, La Fondation de France, ALFEDIAM, Société francophone du diabète, ONIVINS, Abbott, Ardix Medical, Bayer Diagnostics, Becton Dickinson, Cardionics, Merck Santé, Novo Nordisk, Pierre Fabre, Roche, Topcon.

The D.E.S.I.R. Study Group. INSERM U1018: B. Balkau, P. Ducimetière, E. Eschwège; INSERM U1138: F. Alhenc-Gelas; CHU D’Angers: A. Girault; Bichat Hospital: F. Fumeron, M. Marre, R Roussel; CHU de Rennes: F. Bonnet; CNRS UMR8090, Lille: A. Bonnefond, P. Froguel; Université Paris Descartes, UMR1153, Paris: F. Rancière; Centres d’Examens de Santé: Alençon, Angers, Blois, Caen, Chateauroux, Chartres, Cholet, Le Mans, Orléans, Tours; Institute de Recherche Médecine Générale: J. Cogneau; General practitioners of the Region; Institute inter-Regional pour la Santé: C. Born, E. Caces, M. Cailleau, O Lantieri, J.G. Moreau, F. Rakotozafy, J. Tichet, S. Vol.

The COFRASA (clinicalTrial.gov number NCT 00338676) and GENERAC (clinicalTrial.gov number NCT00647088) studies are supported by grants from the Assistance Publique - Hôpitaux de Paris (PHRC National 2005 and 2010, and PHRC regional 2007).

## Author contributions

S.T., C.D., D.M.Z., S.L.S., V.P., R.Re., P.P., P.M., T.L.T., Y.B. and J.J.S. contributed to the conception and study design.

S.T., C.D., D.M.Z., N.G., F.S., M.A.C., B.A., A.S.B., E.B., S.B., I.F., J.C.R., J.P.V, C.B., V.P., F.D., P.P., P.M., T.L.T., Y.B. and J.J.S. contributed to data collection.

S.T., C.D., S.L.S., R.C., S.R., Z.L., F.S., M.L., S.L., M.K., E.C., R.Ro. and Y.B. contributed to data analysis.

S.T., C.D., S.L.S., N.G., P.M., Y.B. and J.J.S. contributed to data interpretation.

S.T. and Y.B. drafted the manuscript.

C.D., D.M.Z., S.L.S., R.C., N.G., S.R., P.P., P.M., T.L.T. and J.J.S. contributed to and revised the manuscript.

## Competing financial interests

All authors declare no competing interests.

## References

1. Freeman, R.V. & Otto, C.M. Spectrum of calcific aortic valve disease: pathogenesis, disease progression, and treatment strategies. Circulation 111, 3316–26 (2005).

2. Rajamannan, N.M., Bonow, R.O. & Rahimtoola, S.H. Calcific aortic stenosis: an update. Nat Clin Pract Cardiovasc Med 4, 254–62 (2007).

3. Chan, K.L., Teo, K., Dumesnil, J.G., Ni, A. & Tam, J. Effect of Lipid lowering with rosuvastatin on progression of aortic stenosis: results of the aortic stenosis progression observation: measuring effects of rosuvastatin (ASTRONOMER) trial. Circulation 121, 306–14 (2010).

4. Rossebo, A.B. et al. Intensive Lipid Lowering with Simvastatin and Ezetimibe in Aortic Stenosis. N Engl J Med 359, 1343–56 (2008).

5. Cowell, S.J. et al. A randomized trial of intensive lipid-lowering therapy in calcific aortic stenosis. N Engl J Med 352, 2389–97 (2005).

6. Martinsson, A. et al. Familial Aggregation of Aortic Valvular Stenosis: A Nationwide Study of Sibling Risk. Circ Cardiovasc Genet 10(2017).

7. Probst, V. et al. Familial aggregation of calcific aortic valve stenosis in the western part of France. Circulation 113, 856–60 (2006).

8. Thériault, S. et al. A transcriptome-wide association study identifies PALMD as a susceptibility gene for calcific aortic valve stenosis. Nat Commun 9, 988 (2018).

9. Thanassoulis, G. et al. Genetic associations with valvular calcification and aortic stenosis. N Engl J Med 368, 503–12 (2013).

10. Helgadottir, A. et al. Genome-wide analysis yields new loci associating with aortic valve stenosis. Nat Commun 9, 987 (2018).

11. McCarthy, S. et al. A reference panel of 64,976 haplotypes for genotype imputation. Nat Genet 48, 1279–83 (2016).

12. Giambartolomei, C. et al. Bayesian test for colocalisation between pairs of genetic association studies using summary statistics. PLoS Genet 10, e1004383 (2014).

13. Kichaev, G. et al. Integrating functional data to prioritize causal variants in statistical fine-mapping studies. PLoS Genet 10, e1004722 (2014).

14. Zhou, J. & Troyanskaya, O.G. Predicting effects of noncoding variants with deep learning-based sequence model. Nat Methods 12, 931–4 (2015).

15. Consortium, G.T. et al. Genetic effects on gene expression across human tissues. Nature 550, 204–213 (2017).

16. El Husseini, D. et al. P2Y2 receptor represses IL-6 expression by valve interstitial cells through Akt: implication for calcific aortic valve disease. J Mol Cell Cardiol 72, 146–56 (2014).

17. Riege, K. et al. Massive Effect on LncRNAs in Human Monocytes During Fungal and Bacterial Infections and in Response to Vitamins A and D. Sci Rep 7, 40598 (2017).

18. Wang, Y. et al. AS-IL6 promotes glioma cell invasion by inducing H3K27Ac enrichment at the IL6 promoter and activating IL6 transcription. FEBS Lett 590, 4586–4593 (2016).

19. Orimo, H. The mechanism of mineralization and the role of alkaline phosphatase in health and disease. J Nippon Med Sch 77, 4–12 (2010).

20. Schlotter, F. et al. Spatiotemporal Multi-omics Mapping Generates a Molecular Atlas of the Aortic Valve and Reveals Networks Driving Disease. Circulation (2018).

21. Verma, S.S. et al. Identifying Genetic Associations with Variability in Metabolic Health and Blood Count Laboratory Values: Diving into the Quantitative Traits by Leveraging Longitudinal Data from an Ehr. Pac Symp Biocomput 22, 533–544 (2017).

22. Chambers, J.C. et al. Genome-wide association study identifies loci influencing concentrations of liver enzymes in plasma. Nat Genet 43, 1131–8 (2011).

23. Kanai, M. et al. Genetic analysis of quantitative traits in the Japanese population links cell types to complex human diseases. Nat Genet 50, 390–400 (2018).

24. Morris, J.A. et al. An atlas of genetic influences on osteoporosis in humans and mice. Nat Genet (2018).

25. Maes, T., Barcelo, A. & Buesa, C. Neuron navigator: a human gene family with homology to unc-53, a cell guidance gene from Caenorhabditis elegans. Genomics 80, 21–30 (2002).

26. Hoffmann, T.J. et al. Genome-wide association analyses using electronic health records identify new loci influencing blood pressure variation. Nat Genet 49, 54–64 (2017).

27. Evangelou, E. et al. Genetic analysis of over 1 million people identifies 535 new loci associated with blood pressure traits. Nat Genet (2018).

28. Capoulade, R. et al. Relationship Between Proximal Aorta Morphology and Progression Rate of Aortic Stenosis. J Am Soc Echocardiogr 31, 561–569 e1 (2018).

29. Rahimi, K. et al. Elevated blood pressure and risk of aortic valve disease: a cohort analysis of 5.4 million UK adults. Eur Heart J 39, 3596–3603 (2018).

30. Pers, T.H. et al. Biological interpretation of genome-wide association studies using predicted gene functions. Nat Commun 6, 5890 (2015).

31. Dunham, I., Kulesha, E., Iotchkova, V., Morganella, S. & Birney, E. FORGE: A tool to discover cell specific enrichments of GWAS associated SNPs in regulatory regions. F1000Research (2015).

32. Iotchkova, V. et al. Discovery and refinement of genetic loci associated with cardiometabolic risk using dense imputation maps. Nat Genet 48, 1303–1312 (2016).

33. Bulik-Sullivan, B.K. et al. LD Score regression distinguishes confounding from polygenicity in genome-wide association studies. Nat Genet 47, 291–5 (2015).

34. Das, S. et al. Next-generation genotype imputation service and methods. Nat Genet 48, 1284–1287 (2016).

35. Marchini, J., Howie, B., Myers, S., McVean, G. & Donnelly, P. A new multipoint method for genome-wide association studies by imputation of genotypes. Nat Genet 39, 906–13 (2007).

36. Bezzina, C.R. et al. Common variants at SCN5A-SCN10A and HEY2 are associated with Brugada syndrome, a rare disease with high risk of sudden cardiac death. Nat Genet 45, 1044–9 (2013).

37. Balkau, B. et al. Predicting diabetes: clinical, biological, and genetic approaches: data from the Epidemiological Study on the Insulin Resistance Syndrome (DESIR). Diabetes Care 31, 2056–61 (2008).

38. Purcell, S. et al. PLINK: a tool set for whole-genome association and population-based linkage analyses. Am J Hum Genet 81, 559–75 (2007).

39. Frayley, C. & Raftery, A.E. MCLUST Version 3 for R: Normal Mixture Modeling and Model-Based Clustering. (Department of Statistics, University of Washington, Seattle, 2006).

40. Byers, S. & Raftery, A.E. Nearest-neighbor clutter removal for estimating features in spatial point processes. J Am Stat Assoc 93, 577–584 (1998).

41. Sudlow, C. et al. UK biobank: an open access resource for identifying the causes of a wide range of complex diseases of middle and old age. PLoS Med 12, e1001779 (2015).

42. Willer, C.J., Li, Y. & Abecasis, G.R. METAL: fast and efficient meta-analysis of genomewide association scans. Bioinformatics 26, 2190–1 (2010).

43. Gusev, A. et al. Integrative approaches for large-scale transcriptome-wide association studies. Nat Genet 48, 245–52 (2016).

44. Finucane, H.K. et al. Partitioning heritability by functional annotation using genome-wide association summary statistics. Nat Genet 47, 1228–35 (2015).

45. Bulik-Sullivan, B. et al. An atlas of genetic correlations across human diseases and traits. Nat Genet 47, 1236–41 (2015).

46. Nikpay, M. et al. A comprehensive 1,000 Genomes-based genome-wide association meta-analysis of coronary artery disease. Nat Genet 47, 1121–1130 (2015).

47. Locke, A.E. et al. Genetic studies of body mass index yield new insights for obesity biology. Nature 518, 197–206 (2015).

48. Shungin, D. et al. New genetic loci link adipose and insulin biology to body fat distribution. Nature 518, 187–196 (2015).

49. Morris, A.P. et al. Large-scale association analysis provides insights into the genetic architecture and pathophysiology of type 2 diabetes. Nat Genet 44, 981–90 (2012).

50. Willer, C.J. et al. Discovery and refinement of loci associated with lipid levels. Nat Genet 45, 1274–1283 (2013).

51. Malik, R. et al. Multiancestry genome-wide association study of 520,000 subjects identifies 32 loci associated with stroke and stroke subtypes. Nat Genet 50, 524–537 (2018).

52. Pattaro, C. et al. Genetic associations at 53 loci highlight cell types and biological pathways relevant for kidney function. Nat Commun 7, 10023 (2016).

