## Supplementary Material for "Genetic association analyses highlight *IL6*, *ALPL*, and *NAV1* as three new susceptibility genes underlying calcific aortic valve stenosis"

**Supplementary Table 1**. Lead SNPs associated with CAVS with $P<1x10^{-5}$ in the meta-analysis

| rsID | Chr | Pos | RA | OA | RAF | Beta | SE | OR | P | Dir | I2 | HetP | Nearest gene(s) |
|---|---|---|---|---|---|---|---|---|---|---|---|---|---|
| rs10455872 | 6 | 161010118 | G | A | 0.081 | 0.429 | 0.046 | 1.54 | 6.48E-21 | ++++ | 37 | 0.19 | LPA |
| rs7543039 | 1 | 100049648 | T | C | 0.514 | 0.215 | 0.026 | 1.24 | 1.24E-16 | ++++ | 0 | 0.63 | PALMD |
| rs2069832 | 7 | 22767433 | A | G | 0.428 | 0.151 | 0.027 | 1.16 | 1.11E-08 | ++++ | 62 | 0.05 | IL6 |
| rs12141569 | 1 | 21873002 | C | T | 0.470 | 0.143 | 0.026 | 1.15 | 3.93E-08 | ++++ | 12 | 0.33 | ALPL |
| rs114314058 | 2 | 150926833 | C | G | 0.018 | 0.482 | 0.095 | 1.62 | 4.43E-07 | ++++ | 32 | 0.22 | MMADHC-RND3 |
| rs113277183 | 9 | 16350313 | C | G | 0.016 | 0.535 | 0.109 | 1.71 | 8.94E-07 | ++++ | 10 | 0.34 | C9orf92-BNC2 |
| rs2150026 | 1 | 170645489 | C | T | 0.290 | 0.142 | 0.029 | 1.15 | 1.01E-06 | ++++ | 57 | 0.07 | PRRX1 |
| rs6715876 | 2 | 65486928 | C | T | 0.361 | 0.130 | 0.027 | 1.14 | 1.11E-06 | ++++ | 17 | 0.31 | ACTR2 |
| rs665770 | 1 | 201748124 | A | G | 0.398 | 0.126 | 0.026 | 1.13 | 1.46E-06 | ++++ | 12 | 0.34 | NAV1 |
| rs17010961 | 4 | 86723103 | A | T | 0.141 | 0.181 | 0.038 | 1.20 | 1.49E-06 | ++++ | 37 | 0.19 | ARHGAP24 |
| rs117206641 | 12 | 133086888 | T | C | 0.108 | 0.203 | 0.042 | 1.22 | 1.52E-06 | ++++ | 0 | 0.72 | FBRSL1 |
| rs11180610 | 12 | 76040394 | T | C | 0.189 | 0.175 | 0.037 | 1.19 | 2.28E-06 | ++++ | 0 | 0.79 | KRR1-PHLDA1 |
| rs4932408 | 15 | 89231801 | A | G | 0.013 | 0.615 | 0.131 | 1.85 | 2.56E-06 | ???+ | 0 | 1.00 | ISG20-ACAN |
| rs11234705 | 11 | 86298700 | C | G | 0.436 | 0.121 | 0.026 | 1.13 | 3.37E-06 | ++++ | 44 | 0.15 | ME3 |
| rs2421649 | 3 | 169197333 | G | A | 0.498 | 0.121 | 0.026 | 1.13 | 3.68E-06 | ++++ | 0 | 0.77 | MECOM |
| rs114795211 | 4 | 97916536 | A | G | 0.027 | 0.392 | 0.085 | 1.48 | 3.71E-06 | ++++ | 9 | 0.35 | PDHA2-STPG2 |
| rs12264978 | 10 | 119483869 | T | C | 0.015 | 0.933 | 0.202 | 2.54 | 4.05E-06 | +++? | 0 | 0.47 | EMX2-RAB11FIP2 |
| rs10124723 | 9 | 19257057 | G | C | 0.429 | 0.121 | 0.026 | 1.13 | 4.08E-06 | ++++ | 43 | 0.15 | DENND4C |
| rs7593336 | 2 | 145836429 | G | A | 0.387 | 0.122 | 0.027 | 1.13 | 4.18E-06 | ++++ | 0 | 0.54 | TEX41 |
| rs73056108 | 3 | 31296043 | A | C | 0.091 | 0.234 | 0.051 | 1.26 | 4.80E-06 | ++++ | 0 | 0.73 | MIR466-STT3B |
| rs7937669 | 11 | 42869801 | T | G | 0.338 | 0.125 | 0.027 | 1.13 | 4.88E-06 | ++++ | 0 | 0.71 | LRRC4C-API5 |
| rs112277963 | 11 | 115856313 | C | T | 0.011 | 0.528 | 0.116 | 1.70 | 4.96E-06 | ?+++ | 0 | 0.88 | CADM1-BUD13 |
| rs7748777 | 6 | 41133806 | A | G | 0.448 | 0.120 | 0.027 | 1.13 | 5.68E-06 | ++++ | 0 | 0.43 | TREM2-TREML2 |
| rs12605367 | 18 | 13686334 | T | C | 0.709 | 0.133 | 0.029 | 1.14 | 6.10E-06 | ++++ | 0 | 0.45 | FAM210A |

| | | | | | | | | | | | | | |
|---|---|---|---|---|---|---|---|---|---|---|---|---|---|
| rs170828 | 1 | 59101134 | G | T | 0.118 | 0.170 | 0.038 | 1.19 | 7.05E-06 | ++++ | 0 | 0.86 | TACSTD2-MYSM1 |
| rs12465180 | 2 | 1326143 | C | G | 0.121 | 0.170 | 0.038 | 1.19 | 8.80E-06 | ++++ | 0 | 0.64 | SNTG2 |

Results were corrected for genomic inflation in individual cohorts and at the meta-analysis stage (double correction). Chr: Chromosome; Pos: Position (hg19); RA: Risk allele; OA: Other allele; RAF: Risk allele frequency (UK Biobank); Beta: Logistic regression coefficient; SE: Standard error of logistic regression coefficient; OR: Odds ratio for the risk allele; P: *P* value; Dir: Direction of effect in the four cohorts (QUEBEC-CAVS, CAVS-France-1, CAVS-France-2, UK Biobank); $I^2$: $I^2$ statistic for heterogeneity; HetP: Heterogeneity *P* value.

**Supplementary Table 2.** Predicted effects on chromatin features for lead GWAS SNPs and plausible causal variants identified using PAINTOR at the three new CAVS loci using the DeepSEA algorithm.

SEE EXCEL FILE

**Supplementary Table 3**. Gene set enrichment using DEPICT (first 20 gene sets)

| **Original gene set ID** | **Original gene set description** | **Nominal *P* value** |
|---|---|---|
| MP:0008735 | Increased susceptibility to endotoxin shock | 6.47E-06 |
| MP:0005592 | Abnormal vascular smooth muscle morphology | 4.23E-05 |
| ENSG00000129991 | TNNI3 subnetwork | 4.88E-05 |
| MP:0004148 | Increased compact bone thickness | 5.52E-05 |
| MP:0001726 | Abnormal allantois morphology | 9.97E-05 |
| MP:0005599 | Increased cardiac muscle contractility | 1.21E-04 |
| MP:0002833 | Increased heart weight | 1.56E-04 |
| MP:0008997 | Increased blood osmolality | 1.94E-04 |
| MP:0002190 | Disorganized myocardium | 2.01E-04 |
| MP:0003352 | Increased circulating renin level | 2.25E-04 |
| ENSG00000095794 | CREM subnetwork | 2.98E-04 |
| ENSG00000092847 | EIF2C1 subnetwork | 4.09E-04 |
| MP:0006143 | Increased systemic arterial diastolic blood pressure | 4.31E-04 |
| ENSG00000168610 | STAT3 subnetwork | 5.04E-04 |
| GO:0010675 | Regulation of cellular carbohydrate metabolic process | 6.45E-04 |
| MP:0004564 | Enlarged myocardial fiber | 6.72E-04 |
| GO:0006109 | Regulation of carbohydrate metabolic process | 7.23E-04 |
| GO:0010906 | Regulation of glucose metabolic process | 7.68E-04 |
| ENSG00000168264 | IRF2BP2 subnetwork | 7.92E-04 |
| ENSG00000181929 | PRKAG1 subnetwork | 8.00E-04 |

**Supplementary Table 4**. Sources of summary statistics from GWAS meta-analysis on cardiovascular traits used in the genetic correlation analyses

| **Source** | **Variable(s) used** | **Sample** | **URL** |
|---|---|---|---|
| CARDIoGRAM[45]plusC4D | CAD, MI | 60,801 CAD cases (approximately 70% MI) and 123,504 controls in predominantly Europeans (77%) | http://www.cardiogramplusc4d.org/ |
| UK Biobank (Neale's group analysis) | SBP, DBP | 317,754 Europeans (SBP)<br>317,756 Europeans (DBP) | http://ldsc.broadinstitute.org/gwashare/ |
| Genetic Investigation of Anthropometric Traits (GIANT)[46,47] | BMI | 322,154 Europeans | https://www.broadinstitute.org/collaboration/giant/index.php/GIANT_consortium_data_files |
| | WHR, WHR adjusted for BMI | 210,088 Europeans | |
| Diabetes Genetics Replication and Meta-Analysis (DIAGRAM)[48] | Type 2 diabetes (T2D) | 12,171 T2D cases and 56,862 controls in Europeans | http://diagram-consortium.org/index.html |
| Global Lipids Genetics Consortium (GLGC)[49] | TC, LDL, HDL, TG | 188,577 Europeans | http://csg.sph.umich.edu//abecasis/public/lipids2013/ |
| MEGASTROKE[50] | Ischemic stroke and subtypes | 34,217 ischemic stroke cases (4,373 LAS; 7,193 CES; 5,386 SVS) and 406,111 controls in Europeans | http://megastroke.org/ |
| CKDGen[51] | CKD | 12,385 cases and 104,780 controls in Europeans | http://ckdgen.imbi.uni-freiburg.de/ |

CAD: Coronary artery disease; MI: Myocardial infarction; SBP: Systolic blood pressure; DBP: Diastolic blood pressure; BMI: Body-mass index; WHR: Waist-to-hip ratio; LDL: Low-density lipoprotein cholesterol; HDL: High-density lipoprotein cholesterol; LAS: Large-artery stroke; CES: Cardioembolic stroke; SVS: Small-vessel stroke; CKD: Chronic kidney disease.

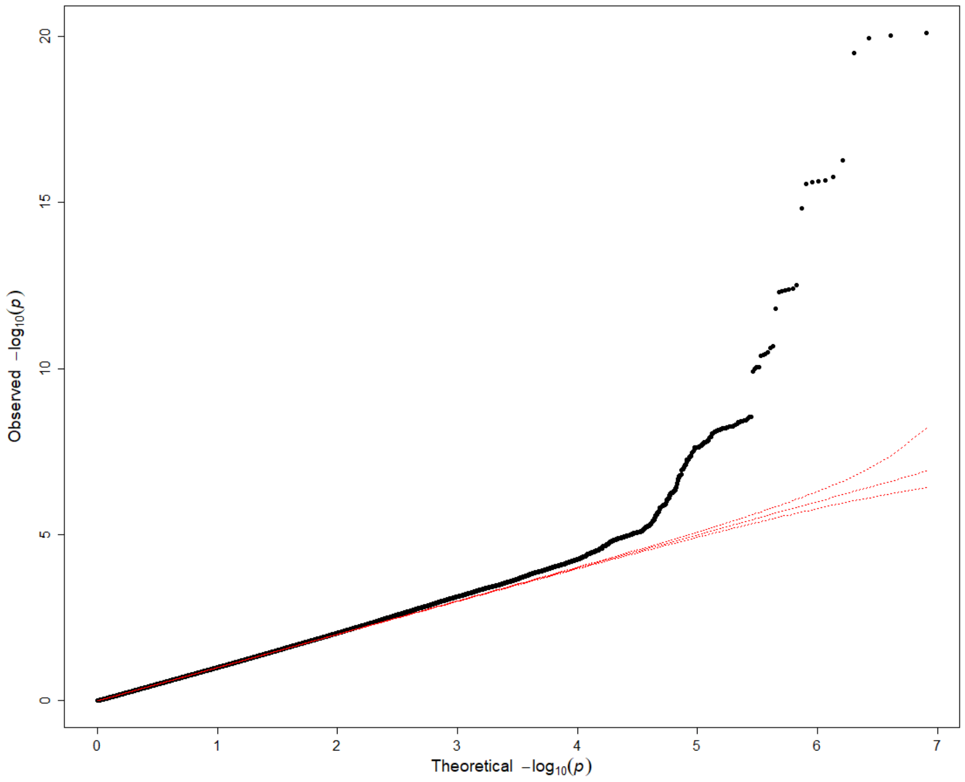


**Supplementary Figure 1**. Quantile-quantile plot of test statistics generated by the GWAS meta-analysis of four cohorts including 5,115 cases and 354,072 controls.

**a** **b**

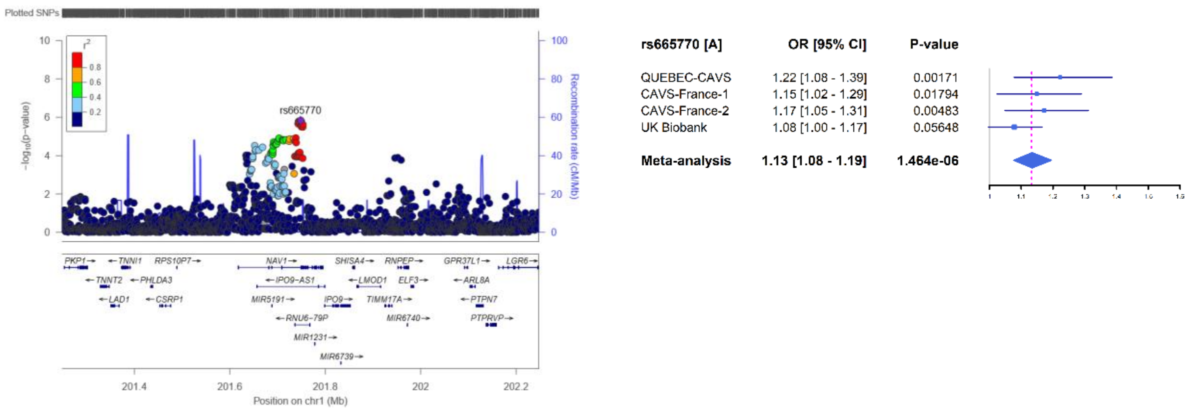


**Supplementary Figure 2**. *NAV1*: a new CAVS-associated loci. (**a**) Regional plot showing the *NAV1* locus. The y axis shows the *P* values in -log10 scale for SNPs up- and downstream of the sentinel SNP (purple dot). The extent of linkage disequilibrium (LD; $r^2$ values) for all SNPs with the sentinel SNP is indicated by colors. The location of genes is shown at the bottom. SNPs are plotted based on their chromosomal position on build hg19. (**b**) Forest plot showing the effect size of the top CAVS-associated SNP at the *NAV1* locus in each cohort and meta-analysis. The blue filled squares represent the odds ratio (OR) for each cohort. The horizontal lines represent the 95% confidence intervals of the OR. The grey and the dashed magenta vertical lines represent an OR of 1.0 and the OR of the meta-analysis, respectively.

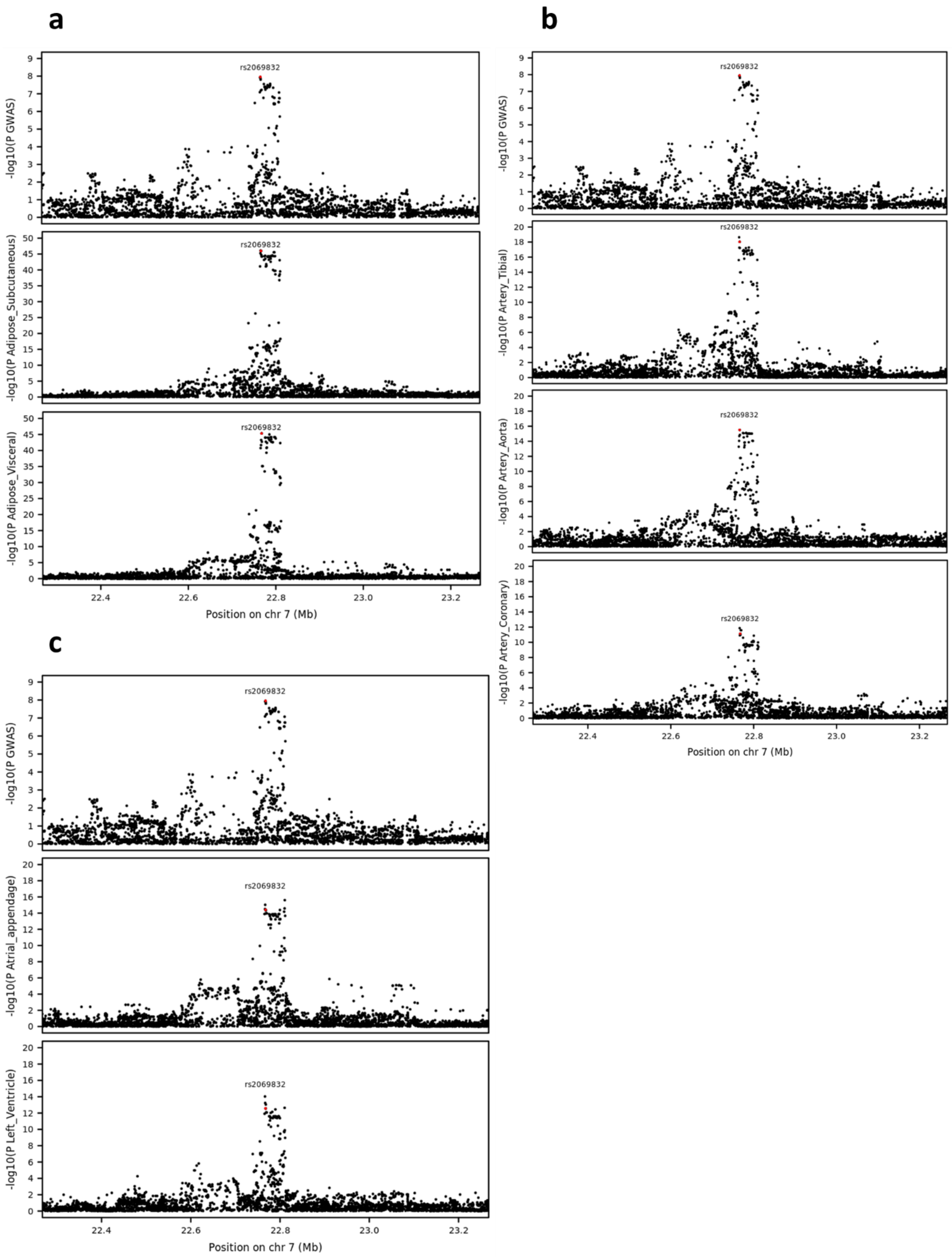
a
b
c
rs2069832
-log10(P GWAS)
-log10(P Adipose_Subcutaneous)
-log10(P Adipose_Visceral)
-log10(P Artery_Tibial)
-log10(P Artery_Aorta)
-log10(P Artery_Coronary)
-log10(P Atrial_appendage)
-log10(P Left_Ventricle)
Position on chr 7 (Mb)
22.4
22.6
22.8
23.0
23.2

**Supplementary Figure 3**. Association with CAVS and the *IL6* RNA antisense (LOC541472) expression in different tissues at the *IL6* locus. (**a**) Adipose tissues. (**b**) Artery tissues. (**c**) Heart tissues. Expression data is from the GTEx project[15]. The GWAS lead SNP, rs2069832, is indicated by a red dot.

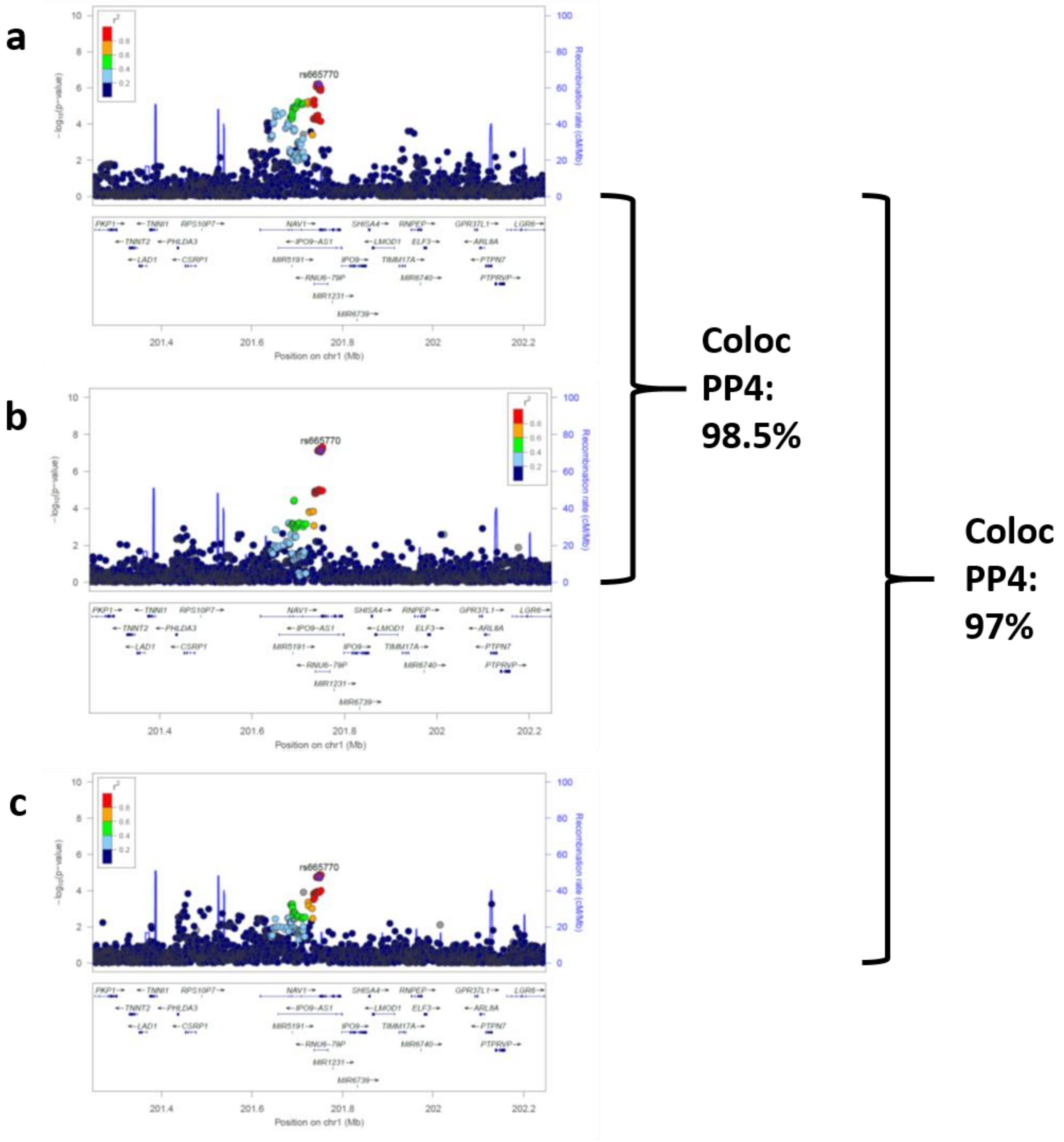


**Supplementary Figure 4**. Colocalization of GWAS signals for the CAVS meta-analysis (**a**) with carotid stenosis (**b**) and carotid procedures (**c**) in UK Biobank.

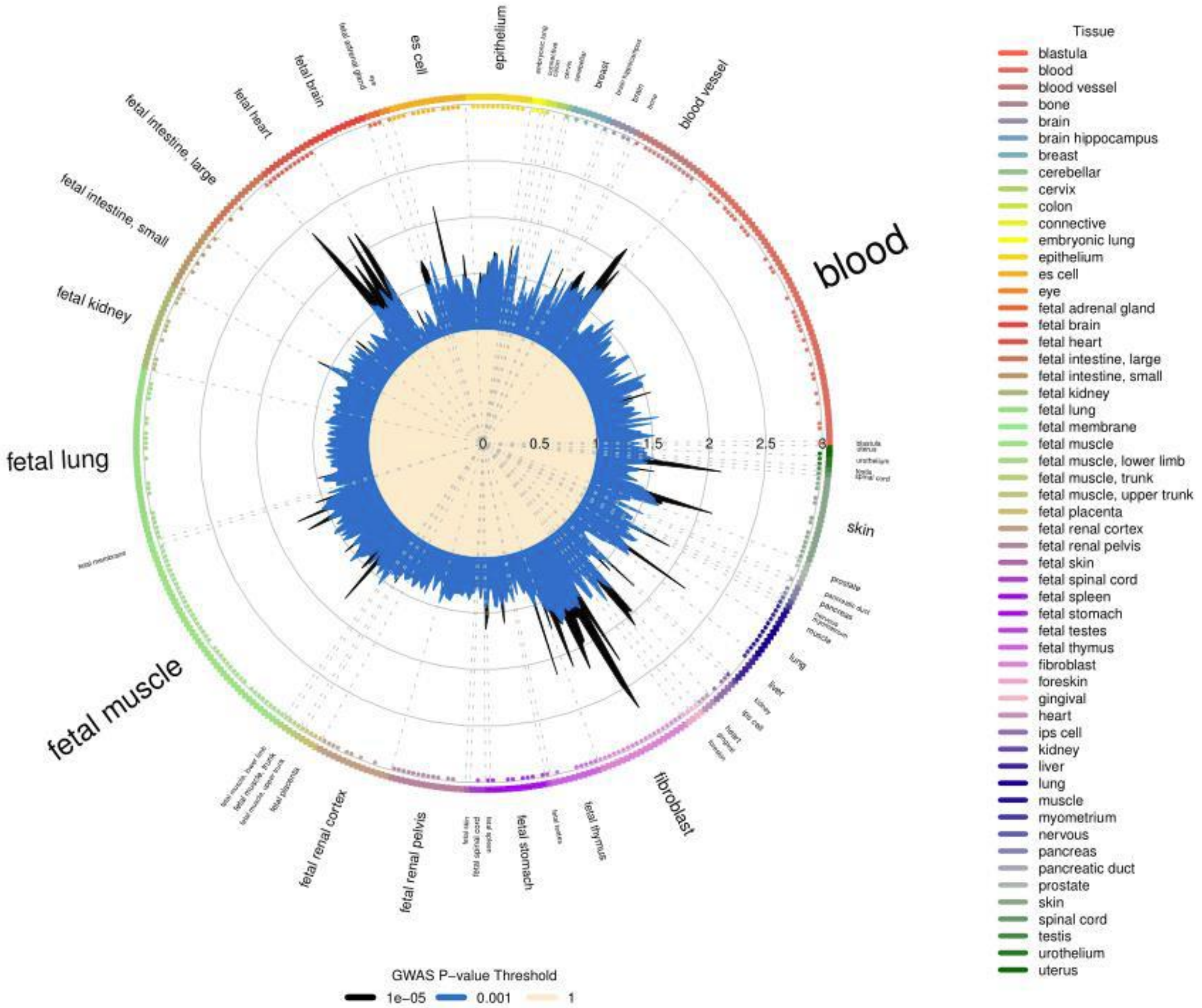


**Supplementary Figure 5**. GARFIELD functional enrichment analyses. Enrichment of CAVS variants in DNase I Hypersensitive sites (broad peaks) from ENCODE and Roadmap Epigenomics data. Radial plot shows the enrichment (OR) in each cell type for two GWAS significance thresholds (0.001 and $1x10^{-5}$).
